# Early endosome disturbance and endolysosomal pathway dysfunction in Duchenne muscular dystrophy

**DOI:** 10.1101/2024.12.16.628552

**Authors:** Julie Chassagne, Nathalie Da Silva, Ines Akrouf, Bruno Cadot, Laura Julien, Ines Barthélémy, Stéphane Blot, Caroline Le Guiner, Mai Thao Bui, Norma B. Romero, Jeanne Lainé, France Pietri-Rouxel, Pierre Meunier, Kamel Mamchaoui, Stéphanie Lorain, Marc Bitoun, Sofia Benkhelifa-Ziyyat

**Author notes:** Correspondence should be addressed to: S. B-Z. 00 33 (0) 1 40 77 81 47. Both authors contributed equally to this work (Julie Chassagne, Nathalie Da Silva).

## Abstract

Duchenne muscular dystrophy (DMD) is a lethal dystrophy characterized by the progressive loss of muscle fibers caused by mutations in *DMD* gene and absence of the dystrophin protein. While autophagy and lysosome biogenesis defects have been described in DMD muscles, the endosomal pathway has never been studied. Here, we showed that impaired lysosome formation is associated with altered acidification and reduced degradative function of the endolysosomal pathway in muscle cells derived from DMD patients. Our data demonstrated that early endosomes are increased in these cells as well as in muscle biopsies from DMD patients and two animal models of DMD, *mdx* mice and GRMD dogs. We determined that these abnormalities are due to the lack of dystrophin *per s*e and could be correlated with disease progression and severity. We further identified an abnormal upregulation of the Rab5 GTPase protein, one key actor of early endosomal biogenesis and fusion, in the three DMD models which may underlie the endosomal defects. Finally, we demonstrated that Rab5 knock-down in human DMD muscle cells as well as dystrophin restoration in GRMD dogs, normalize Rab5 expression levels and rescue endosomal abnormalities. This study unveils a defect in a pathway essential for muscle homeostasis and for efficacy of adeno associated virus vectors and antisense oligonucleotides-mediated therapies.

## Introduction

Duchenne muscular dystrophy (DMD) is the most frequent X-linked lethal form of muscular dystrophies in humans, caused by mutations in the *DMD* gene leading to complete loss of the subsarcolemmal protein dystrophin^1^. Dystrophin is part of the multiprotein dystrophin-associated glycoprotein complex ensuring a link between the extracellular matrix and the cytoskeletal actin that is crucial for the fiber stability and plasma membrane flexibility during muscle contraction. In muscles of DMD patients, lack of dystrophin leads to sarcolemma fragility, frequent cycles of necrosis/regeneration associated with muscle weakness and loss leading to premature death^1^. The majority of DMD patients (∼60-70%) have a deletion in the *DMD* gene affecting one or more exons concentrated between exons 45 and 55. In some cases, *DMD* mutations induce expression of truncated but partially functional dystrophins leading to a milder phenotype known as Becker muscular dystrophy (BMD). The large size of dystrophin’s cDNA makes it difficult to develop therapeutic approaches that deliver the full-length dystrophin. Consequently, two main gene therapies allowing expression of BMD-like dystrophins have been developed: the transfer of cDNAs expressing microdystrophins in muscles using adeno-associated virus (AAV) vectors, and the targeted exon skipping strategy to skip mutated exons and maintain open reading frame using antisense oligonucleotides (ASOs). The latter approach uses among others, 2’OMePS and PMO oligonucleotide chemistries or AAVs for ASO expression. These two approaches have shown promising results in DMD animal models, i.e. *mdx* mouse and golden retriever muscular dystrophy (GRMD) dogs, and have reached clinical trials^2–4^. Noteworthy their therapeutic efficacy depends on disease progression. Indeed, we previously demonstrated that muscle regeneration occurring in the dystrophic context impacts AAV-mediated expression and the subsequent therapeutic benefits in *mdx* mice^5–7^ and that transient dystrophin restoration by an ASO pre-treatment allowed a ten-fold increase of the AAV-mediated dystrophin expression^5^. Cellular and molecular mechanisms disturbing the DMD muscle structure and function, need to be elucidated in order to improve current treatments and develop new therapeutic approaches.

Several dysfunctional pathways have been described in muscles of human and animal DMD models including disorganization of microtubule network^8^ and Golgi complex^9^ as well as increased excretion of exosomes^10^ in *mdx* myofibers. Moreover, the disturbance of Ca^2+^ homeostasis and increased oxidative stress in DMD models were linked to impaired lysosomal biogenesis and autophagy as widely described in several muscular dystrophies, lysosomal storage diseases as well as Down syndrome (DS) and Alzheimer diseases^11–15^. Lysosomes are organelles of low pH (6.5 to 4.5) containing acidic hydrolases required for a proper degradation of the autophagy and endosomal pathway contents^16^. The endosomal pathway contributes to autophagy homeostasis^17,18^ and *vice versa* functional autophagy pathway is required for the optimal endosome function^19^. During endocytosis, vesicles cleaved from the plasma membrane fuse with early endosomes (EEs), where contents (receptors, receptor-ligands…) are sorted and distributed back to the plasma membrane *via* recycling endosomes, to the trans-Golgi network (TGN), or to late endosomes that fuse with lysosomes to form endolysosomes for degradation^20,21^. Early, late and recycling endosomes as well as lysosomes are different stages of a continuous maturation process during which these vesicles undergo fusion and fission reactions determining the endosome number and volume^22,23^. This process is orchestrated by small GTPases of the Rab family and their effectors. Rab5 with its effectors, such as EEA1, regulates both EEs morphology and fusion^24,25^ whereas Rab7 is required for the initiation and maturation of early-to-late endosomes^26,27^. Otherwise, maturation of the endolysosomal pathway depends on the trans-Golgi network which delivers hydrolases to the acidic late endosomes *via* the cation-independent Mannose-6-Phosphate receptor (CI-MPR)^28^. In addition, overexpression or overactivation of Rab5 accelerate endocytosis and leads to aberrant fusion and enlargement of endosomes^18,25,29–32^ that can be rescued by normalizing Rab5 activity^18,32^. It is noteworthy that endosomal pathway integrity is essential for intracellular trafficking of AAVs and ASOs and efficient therapeutics delivery in nuclei^33,34^.

Here, we investigated the endolysosomal pathway in muscle cells from DMD patients and demonstrated that the degradation function as well as acidification of the endolysosomal pathway are affected. We further showed a dysregulation of early endosomes in muscles of DMD patients, *mdx* mice and GRMD dogs that could be exacerbated by the disease progression. This may be due to an abnormal upregulation of the Rab5 protein, the main regulator of the endosomal biogenesis and fusion, in the three DMD models. Finally, we demonstrated that dystrophin restoration in GRMD dogs normalizes Rab5 expression levels and the endocytic pathway.

## Results

### Endolysosomal pathway is defective in muscle cells from DMD patients

Recent studies showed that dystrophin absence disturbs several intracellular pathways in DMD mouse and human models including DMD proliferating myoblasts^12,35,36^. In this context, we investigated the endolysosomal pathway in DMD patient-derived immortalized muscle cell lines. Absence of dystrophin expression in DMD proliferating cells (myoblasts) and in differentiated cells (myotubes) was confirmed at mRNA and protein level (Supplementary Fig. S1), making this cell model valuable for *in vitro* studies. The proteolytic degradation in the endolysosomal pathway was assessed in epidermal growth factor (EGF) starved DMDΔ45-52 myoblasts by following extinction of fluorescence of AlexaFluor-488 tagged EGF during its trafficking from early endosomes to lysosomes for degradation^37^. The mean fluorescence intensity (MFI) of Alexa-488-EGF analyzed by live imaging from 0 to 60 min post-incubation showed that only 10% of total Alexa-488-EGF were present in control cells after 10 min of incubation, whereas about 40% were still detected in DMD cells (Fig. 1A). This significant delay in EGF degradation was observed over the time course in DMD cells when compared to control cells. To assess if the observed proteolysis defect might be due to impaired acidification of the endolysosomal pathway, we analyzed the staining associated with pHrodo-EGF used as pH probe which fluoresces brightly as pH is reduced. The pHrodo-EGF MFI was about 60% lower in DMD myoblasts compared to controls from 2 to 30 min post-incubation (Fig. 1B). After 20 min, the pHrodo-EGF fluorescence was almost not visible in DMD cells whereas the Alexa-488-EGF was still present consistent with an acidification defect of the EGF-containing vesicles. Altogether, these results show an alteration in the degradative capacity associated with an altered acidification of the endolysosomal pathway in DMD cells.

**Figure 1.**
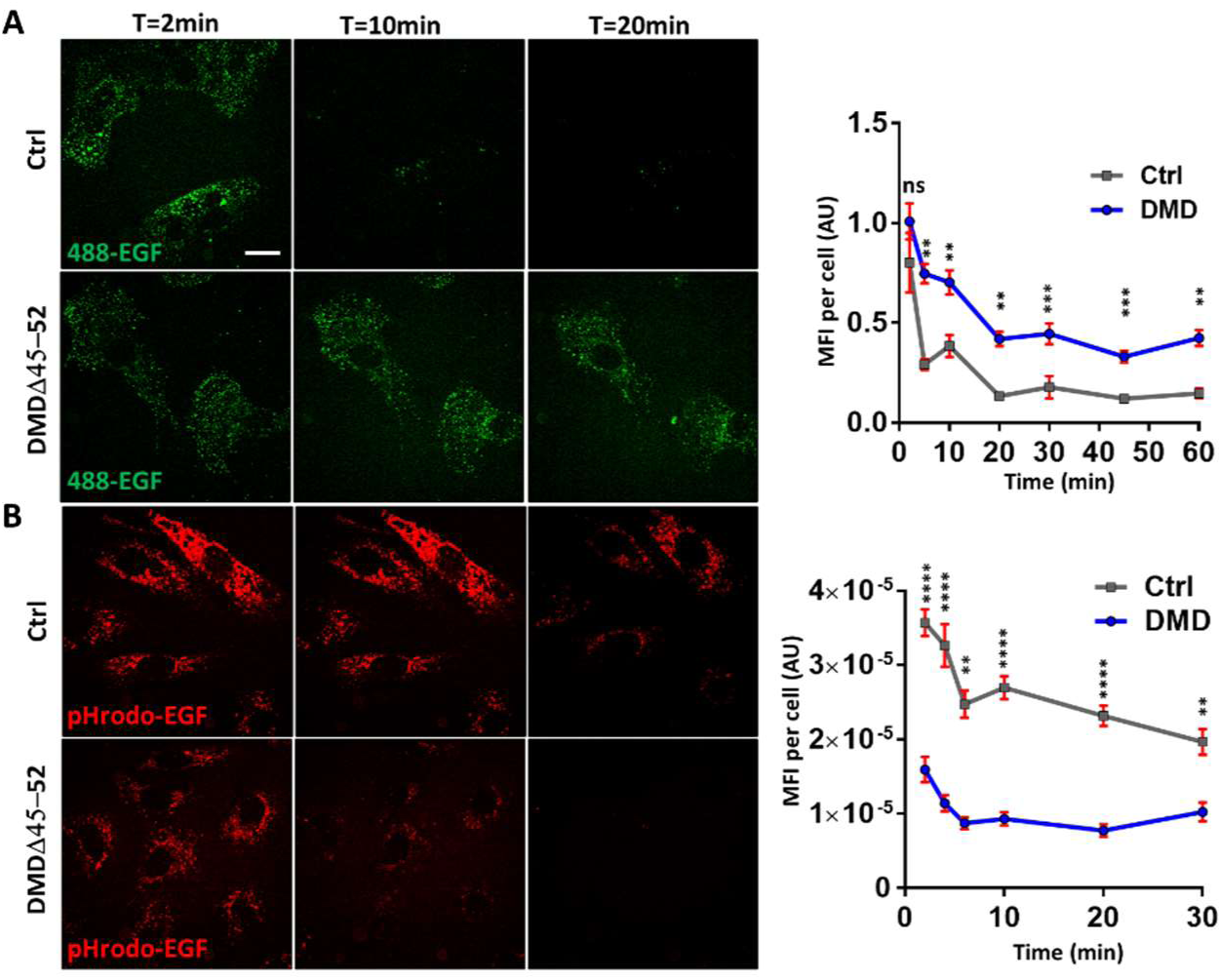
EGF degradation and vesicle acidification in human DMD myoblasts. Live cell imaging of unfixed EGF-starved control (Ctrl) and DMD myoblasts incubated with **(A)** Alexa-Fluor-488-EGF and **(B)** pHrodo-EGF showing a delay in Alexa-488-EGF degradation over the time course in DMD cells compared to control cells and an acidification defect of the pHrodo-EGF-containing vesicles. MFI (mean fluorescence intensity) normalyzed by cell area (µm^2^); AU: arbitrary unit. Scale bar = 20µm. Statistics: one-way ANOVA with a post hoc Bonferroni test. **p<0.01, ***p<0.001, ****p<0.0001, ns: non-significant.

### The early endosomes are dysregulated in muscle cells from DMD patients

To determine the cause of the defect in function of the endolysosomal pathway, we further analyzed the distribution of early endosomes (EE) by immunostainings using the EEs marker EEA1 in immortalized myoblasts and myotubes from two DMD patients (DMDΔ45-52 and DMDΔ52) and two healthy controls (Ctrl1 and Ctrl2). Representative confocal images suggested that EEA1 vesicles localization was mostly similar in DMD and control cells, while their number and size increased both in myoblasts and myotubes of DMD patients compared to controls (Fig. 2A and 2D). Morphometric quantification of EEs carried out on confocal images showed that the numbers of EEA1 positive puncta were between 1.2 and 1.7-fold higher in DMD myoblasts and myotubes compared to controls cells (Fig. 2B and 2E). To ensure that this defect is not due to cell immortalization, similar experiments were performed in native primary myoblasts derived from three DMD patients (DMDΔ45-52, DMDΔ48-50 and DMDΔ44-50) and two healthy controls. In accordance with data in immortalized myoblasts, results showed a higher endosome content in all primary DMD cell lines (Supplementary Fig. S2A) validating the use of the immortalized cells for the rest of our study. To assess whether EE defect is specific of DMD pathophysiological context, morphometric analysis was performed in muscle cells from patients affected by two other muscular dystrophies, Limb-Girdle Muscular Dystrophy type 2C (LGMD2C) and facioscapulohumeral muscular dystrophy type 1 (FSHD1). No or minor change was observed in the number of EEA1 positive puncta in myoblasts derived from these two dystrophies compared to 2 control cell lines (Supplementary Fig. S2B).

**Figure 2.**
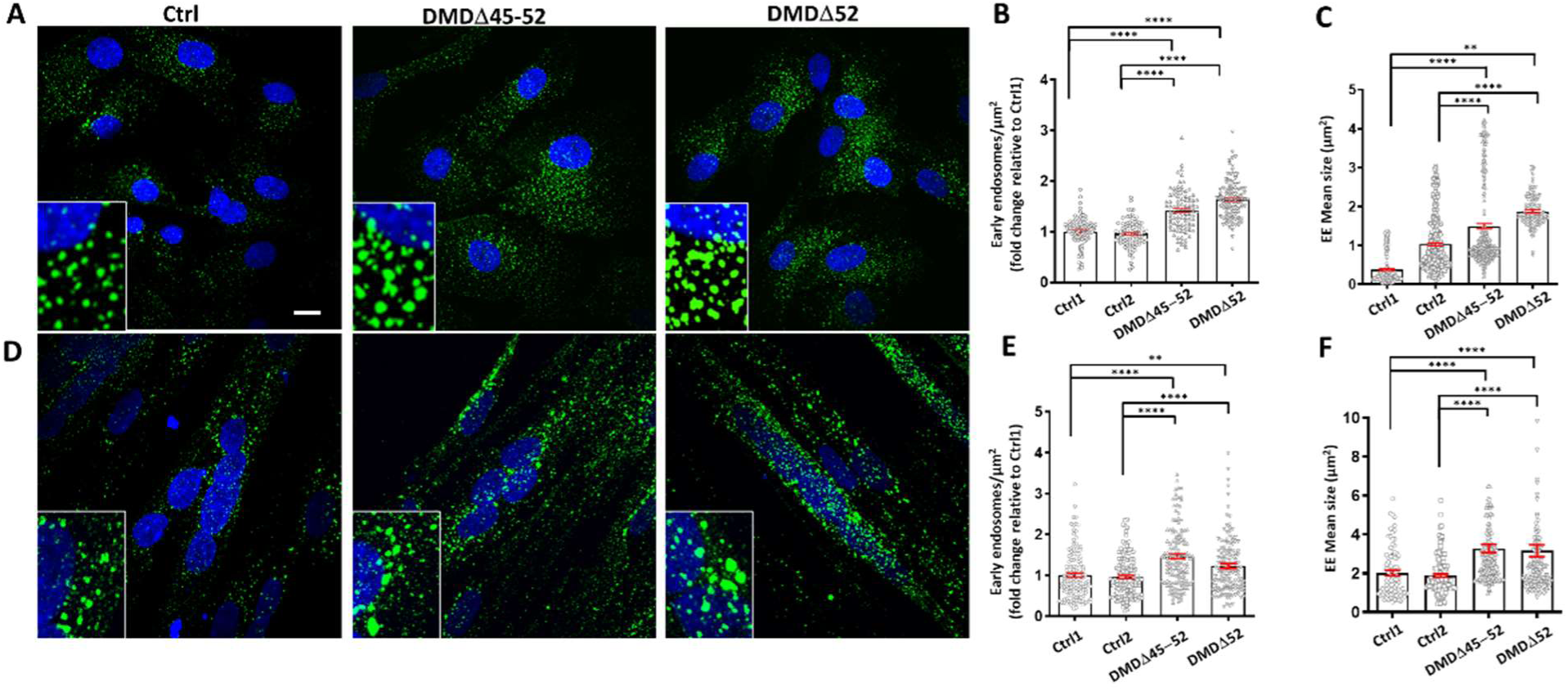
Early endosomes in human DMD myoblasts and myotubes. Representative confocal images of immortalized primary DMDΔ45-52, DMDΔ52 and healthy controls (Ctrl) **(A)** myoblasts and **(D)** myotubes stained with DAPI to mark nuclei (blue) and anti-EEA1 antibody to mark early endosomes (green). Quantification of EEA1 positive puncta normalized by cell area on confocal microscopy images showed an increased number of EEA1 positive puncta in DMD myoblasts **(B)** and myotubes **(E)** compared to Ctrl. The data are represented as the mean ± SEM of EEA1 puncta number per cell area of at least 88 cells per condition (n = 3 independent experiments). Quantification of EEA1 puncta mean size (µm^2^) from confocal microscopy images of **(C)** DMD myoblasts and (**F)** myotubes compared to their respective Ctrl. The data are represented as the mean ± SEM of EEA1 puncta size per cell of at least 100 cells per condition (n = 3 independent experiments). Scale bar = 10µm. Statistics: one-way ANOVA with a post hoc Bonferroni test, **p<0.01, ****p<0.0001.

Our data also revealed that the mean area of EEA1-positive puncta was significantly higher in DMDΔ45-52 and DMDΔ52 myoblasts and myotubes (Fig. 2C and 2F). Electron microscopy (EM) analysis of DMD myoblasts revealed that the increased EEA1-positive staining highlighted by confocal imaging corresponded to both enlarged EEs and grouped EEs (Fig. 3).

**Figure 3.**
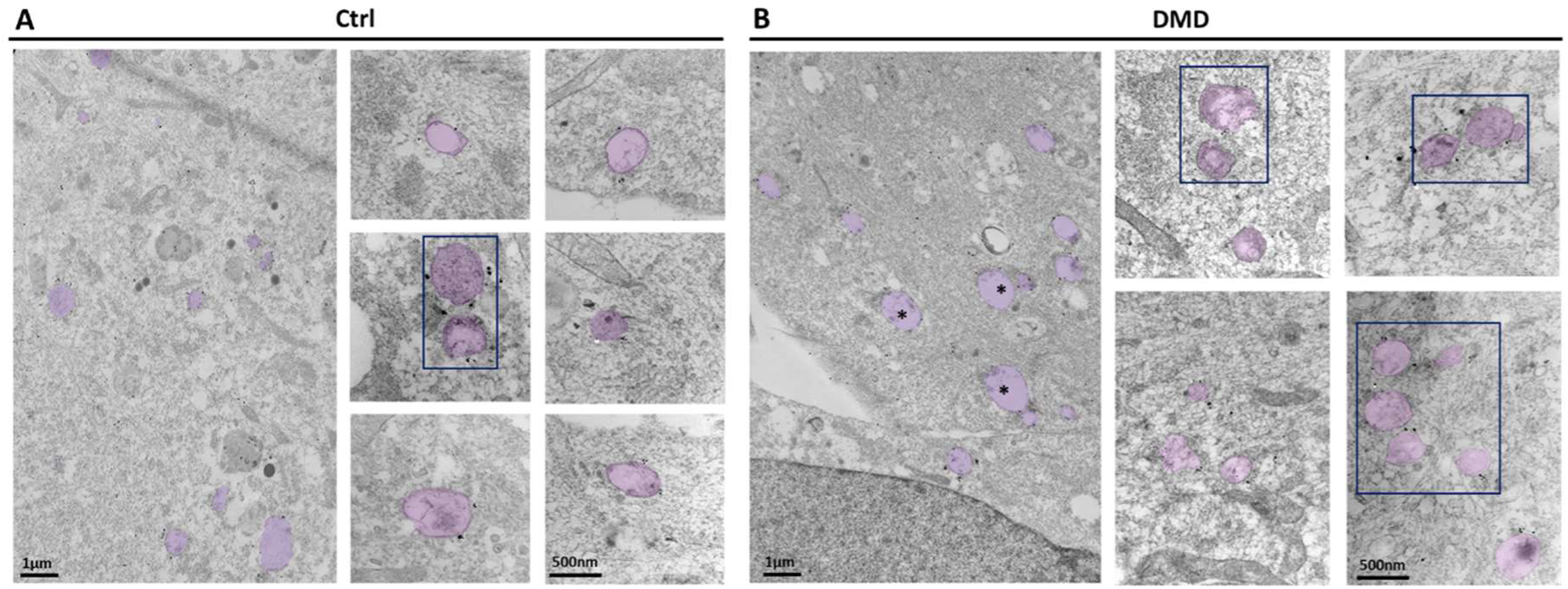
Ultrastructure of early endosomes in myoblasts of DMD patients. Low and high magnification electron micrographs of early endosomes (EEs) in myoblasts from **(A)** healthy controls (Ctrl) and **(B)** DMDΔ45-52 patients after EEA1 immunogold-labeling showing large isolated EEs (> 1µm, asterisk) and clusters (rectangular outlines) of at least 2 endosomes that are more frequently observed in DMD myoblasts compared to controls (Ctrl). Early endosomes are colored in purple for clarity.

To further investigate the endolysosomal abnormalities, we analyzed late endosomes in myoblasts of two DMD patients (DMDΔ45-52 and DMDΔ52) compared to two healthy control cell lines, using the late endosomal marker Rab7 in immunostaining experiment. Results showed that the mean fluorescence intensity (MFI) associated with Rab7 staining was reduced in DMDΔ45-52 (by ̴35%) and DMDΔ52 (by ̴60 %) myoblasts compared to healthy controls (Fig. 4A and 4B). As late endosomes are considered the gateway to lysosomal degradation, we examined previously reported lysosomal defects in *mdx* muscles^12^ in these myoblasts. Consistently, the lysosomal marker LAMP1 staining was reduced in DMDΔ45-52 (−25 to 30%) and DMDΔ52 (−30 to 35%) cells compared to healthy controls (Fig 4C and 4D). In addition, we observed that the large perinuclear lysosomes were missing whereas smaller dots were still present. Altogether, our results showed an endolysosomal dysfunction in muscle cells of DMD patients associated with enlarged and clustered early endosomes as well as a reduction of late endosomes and lysosomes.

**Figure 4.**
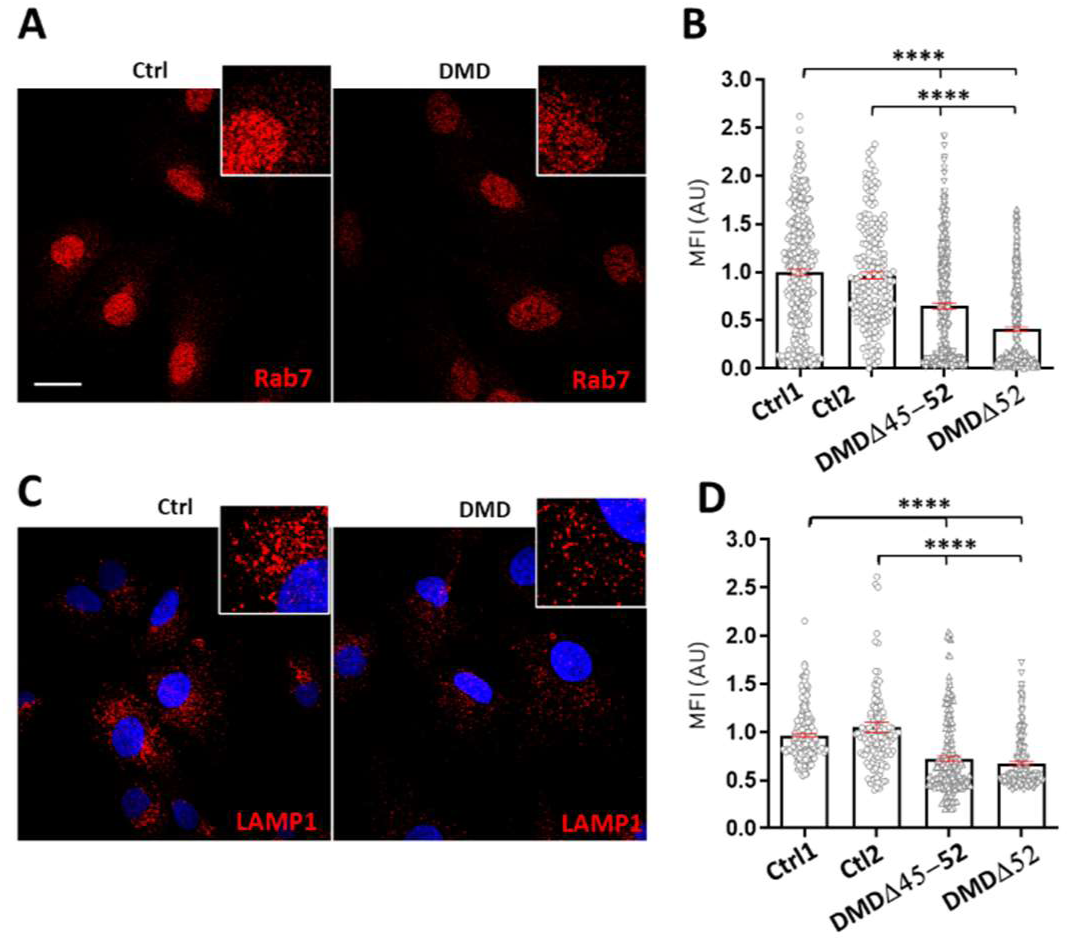
Late endosomes and lysosomes in human DMD myoblasts. Representative confocal images of immortalized primary DMD and healthy control (Ctrl) myoblasts stained with **(A)** anti-Rab7 to mark late endosomes (red) and, **(C)** DAPI to mark nuclei (blue) and anti-Lamp1 antibody to mark lysosomes (red). Quantification of the mean fluorescence intensity (MFI) normalized by cell area of the **(B)** Rab7 and **(D)** LAMP1 immunostainings showed a significant decrease of lysosomes and late endosomes vesicles in DMD myoblasts compared to Ctrl. Scale bar = 10µm. MFI: mean fluorescence intensity; AU: arbitrary unit. Statistics: one-way ANOVA with a post hoc Bonferroni test, ****p<0.0001.

### The early endosomes are disturbed in muscles of DMD patients and two animal models

Early endosomes were analyzed in muscle biopsies from 3-, 5- and 10-year-old DMD patients (respectively DMDΔ45-52, DMDΔ52 and DMDΔ48-50) and GRMD dogs (splice site mutation in Dmd intron 6) as well as in isolated myofibers from *mdx* mice (nonsense mutation in Dmd exon 23). Muscle sections of the three DMD patients showed a higher EEs immunostaining (Fig. 5A) not homogeneously distributed in the fibres when compared with healthy controls. Higher EEs immunostaining was also observed in muscle biopsies of 2 subgroups of GRMD dogs which are severely (loss of ambulation before the age of 6-months) or moderately affected (still ambulant at 6-months) (Fig. 5B and 5C). Interestingly, this increase was more pronounced in the severely affected subgroup, being already statistically significant at an early timepoint (2 months of age) (Fig. 5D and Supplementary Fig. S3A), suggesting that this abnormality is correlated with the disease severity. In the moderately affected subgroup, this increase became significant at 6 months of age, correlating with the disease progression (Fig. 5C and 5D and Supplementary Fig. S3A). In mdx, the analysis was performed in isolated myofibers from 2- and 12-week-old mice i.e. before and after the massive necrosis/regeneration cycles, respectively. As expected, mdx fibers of 12-week-old mice showed centrally located nuclei, a hallmark of regenerating fibers^38^, and a disrupted microtubule cytoskeleton^8,39^ (Fig. 5E). While EEs aligned following the microtubule cytoskeleton of nuclei of the wild-type (WT) muscle fiber, they were non-uniformly distributed in the 12-week-old mdx fibers (Fig. 5E). The difference in EE organization was already measurable in 2-week-old mdx fibers when compared to controls (Supplementary Fig. S3B). As in muscle fibers of DMD patients and GRMD dogs, quantification of EE immunostaining revealed about a 2.2-fold and 2-fold higher number of EE1A positive puncta respectively in 2-week-old and 12-week-old mdx mice compared to age-matched WT mice (Fig. 5F and Fig. 5G).

**Figure 5.**
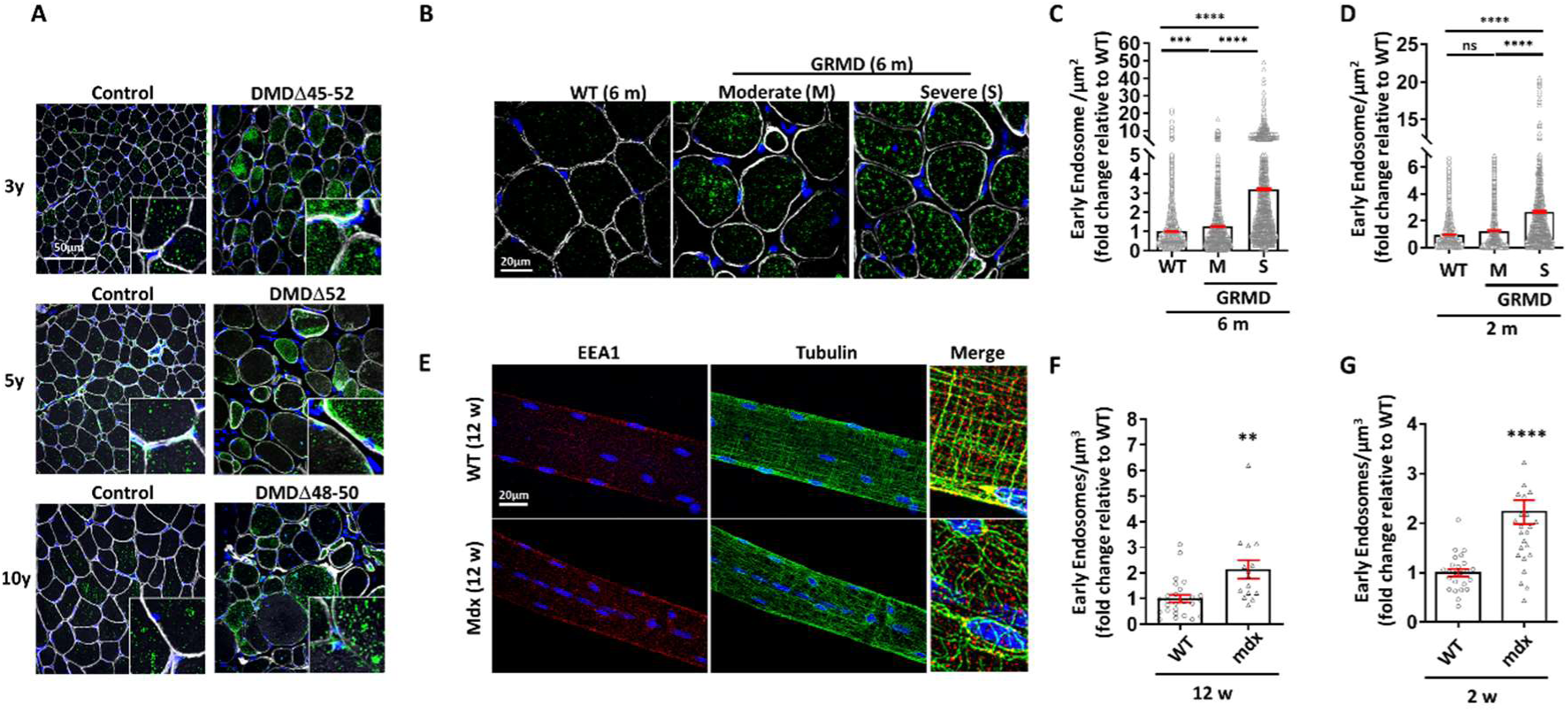
Distribution and quantification of early endosomes in human and GRMD biopsies, and mdx muscle fibers. Representative confocal images showing sections of muscle biopsies from **(A)** 3-, 5- and 10-year-old DMD patients (respectively DMDΔ45-52, DMDΔ52, DMDΔ48-50) and age-matched healthy controls (n=1 for each) and **(B)** severely (S) (n=3) or moderately (M) (n=3) affected GRMD dogs of 6 month-old (6 m) and age-matched healthy control dogs (n=3) stained with DAPI to mark nuclei, anti-caveolin (for dog) or anti-laminin alpha-2 (for human) antibodies to mark the plasma membrane (white) and EEA1 antibody to mark early endosomes (green) or **(E)** myofibers isolated from the EDL muscle of wild type (WT) and mdx of 12 week-old (n=4) labeled with DAPI to mark nuclei (blue), anti-tubulin antibody to mark the microtubule cytoskeleton (green), and anti-EEA1 antibody to mark EEs (red). Quantification of EEA1 positive puncta showed a significant increase in early endosome staining in muscles of **(C)** GRMD of 6-month-old (6 m, n=3) and **(D)** 2-month-old (2 m, n=3) and, **(F)** 12 week-old (12 w, n=4) and **(G)** 2 week-old (2 w, n=3) mdx fibers compared to their respective controls. The data are represented as the mean ± SEM of at least 500 fibers per dog and 15 to 26 myofibers per group of mice analyzed on confocal images. Statistics: one-way ANOVA with a post hoc Bonferroni test, for human and dog analysis and unpaired Student’s t-test (two tailed) for mice. **p<0.01, ***p<0.001, ****p<0.0001, ns: non-significant.

In accordance with our *in vitro* data in DMD patient-derived muscle cells, early endosomes are altered in muscles from DMD patients, GRMD dogs and mdx muscles.

### Rab5 overexpression drives the endosomal abnormalities in DMD models

We examined the expression of Rab5 GTPase, the main regulator of biogenesis and fusion of early endosomes^25,30^, in DMD muscle cells derived from two patients (DMDΔ45-52 and DMDΔ52) (Fig. 6A), in muscle biopsies from GRMD dogs (Fig. 6B) and in muscle from mdx (Fig. 6C) by RT-qPCR and western blot. We showed that Rab5 mRNA and protein were overexpressed (at least 1.8-fold for mRNA) in DMD muscle cells of the three models compared to their respective controls (Fig. 6). To explore whether Rab5 overexpression was responsible for the increased number of EEs, we quantified EEA1 positives puncta in DMDΔ45-52 myoblasts treated with small interfering RNA (siRNA) directed against Rab5 (siRab5) or with a scrambled siRNA negative control (siScr). Two-day treatment of cells with siRNA strongly reduced the Rab5 mRNA (Fig. S4) and normalized Rab5 protein content (Fig. 6D). In this condition, the number of EEA1 positives puncta were rescued in siRab5 treated DMD myoblasts, to a level equivalent to that of control cells (Fig. 6E). To evaluate if the overexpressed Rab5 impacted endocytosis in DMDΔ45-52 myoblasts, we quantified uptake of tagged transferrin (Alexa-488-transferrin: Alexa-488-Tfn or biotin-transferrin: Biot-Tfn), a marker of clathrin-mediated endocytosis internalized and quickly directed to the recycling pathway. Compared to controls, 5- and 10-min uptake of Alexa-488-Tfn and Biot-Tfn analyzed respectively by live imaging and western blot was reduced by 2-fold in DMD cells and achieved normal value 15 min post incubation (Supplementary Fig. S5A and S5B). Since this result correlated with a 2-fold decreased expression of the Tfn receptor observed in DMD cells compared to controls (Supplementary Fig. S5C), these data suggested that the endocytosis was not affected by Rab5 overexpression in DMD cells. Altogether, our data demonstrated a Rab5 overexpression in DMD muscle cells from patients and animal models which could lead to EEs accumulation and swelling.

**Figure 6.**
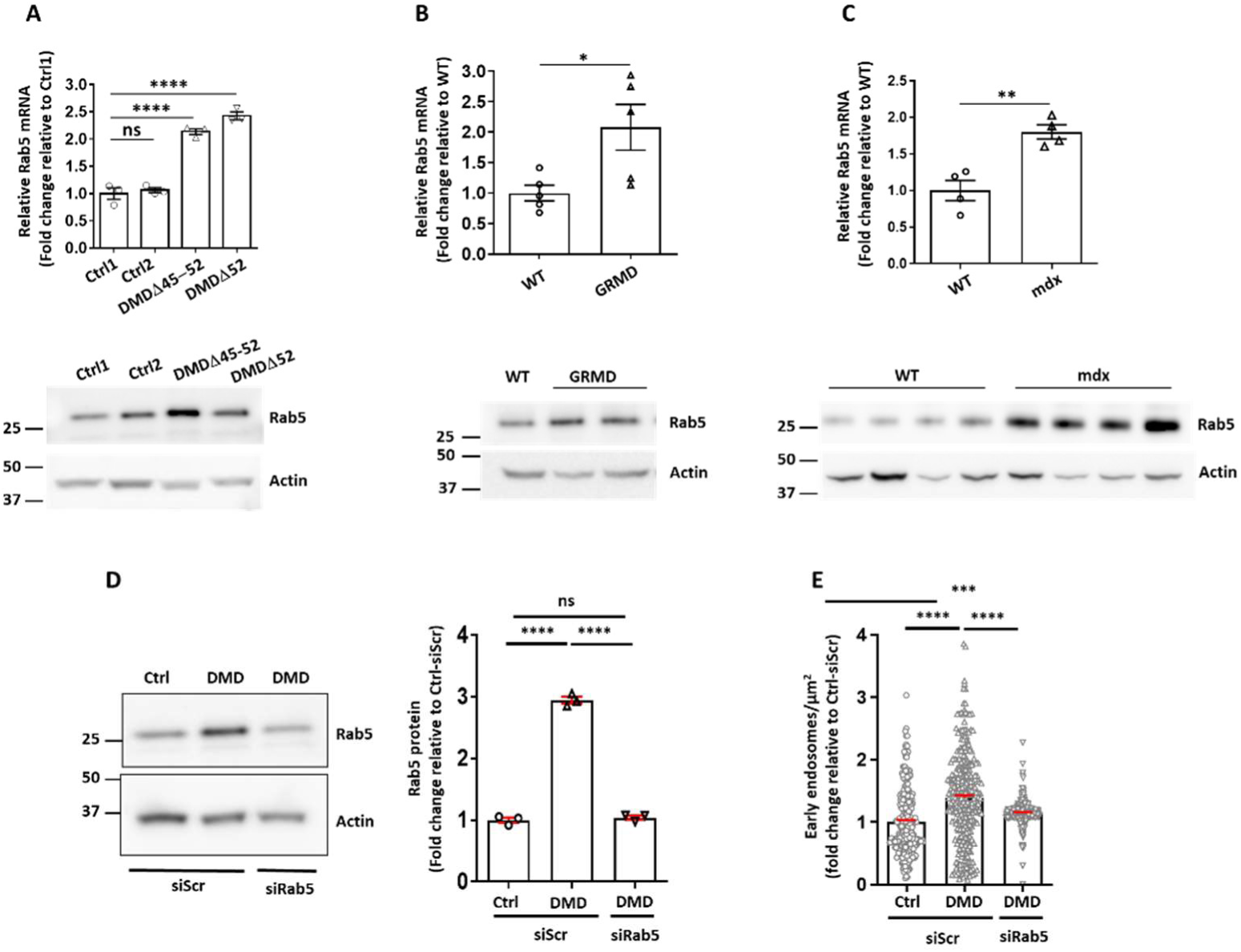
Rab5 expression in DMD models and endosome abnormalities rescue by siRNA-mediated knockdown of Rab5 in DMD myoblasts. The expression of Rab5 mRNA analyzed by RT-PCR (top) and Rab5 protein analyzed by western-blot (bottom) in **(A)** DMD myoblasts (n = 3 independent experiments) **(B)** dog muscles (n = 5 biopsies from 5 WT dogs and n=5 biopsies from 4 GRMD dogs) and **(C)** mdx mouse muscles (n = 4 mice per group). RT-qPCR were performed in duplicate. The data are represented as the mean ± SEM. Statistics: one-way ANOVA with a post hoc Bonferroni test for (A) DMD myoblasts comparison and unpaired Student t-test (two tailed) for (B) dog and (C) mice comparisons. **(D)** Western-blot analysis and quantification (n=3) of Rab5 protein showing the efficacy of Rab5 silencing in human DMDΔ45-52 myoblasts transfected with siRNA directed against Rab5 (siRab5) compared to DMD and control myoblasts (Ctrl) treated with a scrambled siRNA control (siScr). **(E)** The number of EEA1 positive endosomes is reduced in human DMD cells treated with siRab5, to a level equivalent to that of control cells (Ctrl). The data are represented as the mean ± SEM of 3 independent experiments for western-blot and at least 300 cells of three independent experiments analyzed on confocal images for early endosome counting. Statistics: one-way ANOVA with a post hoc Bonferroni test. *p<0.05, **p<0.01, ***p<0.01, ****p<0.0001. ns: non-significant.

### Restoration of dystrophin in GRMD dogs rescued Rab5 overexpression and EE disturbance

To further demonstrate the direct link between absence of dystrophin and endosome disturbance, EEs were analyzed in GRMD dogs treated to rescue dystrophin expression. A one-shot treatment of AAV-U7snRNA vector was shown to be sufficient to elicit substantial levels of restored dystrophin associated with a remarkable improvement of the muscle force in the GRMD dogs^40,41^. We exploited muscle samples of these two studies and used untreated and AAV-U7snRNA treated GRMD dogs well-characterized for dystrophin rescue, to analyze Rab5 expression and EEs. Results demonstrated that the expression of Rab5 mRNA and protein as well as the number of EEs were restored close to WT values in AAV-U7-treated GRMD muscles (Fig. 7A and 7B) revealing that proper Rab5 expression and early endosomes are under the control of the dystrophin expression.

**Figure 7.**
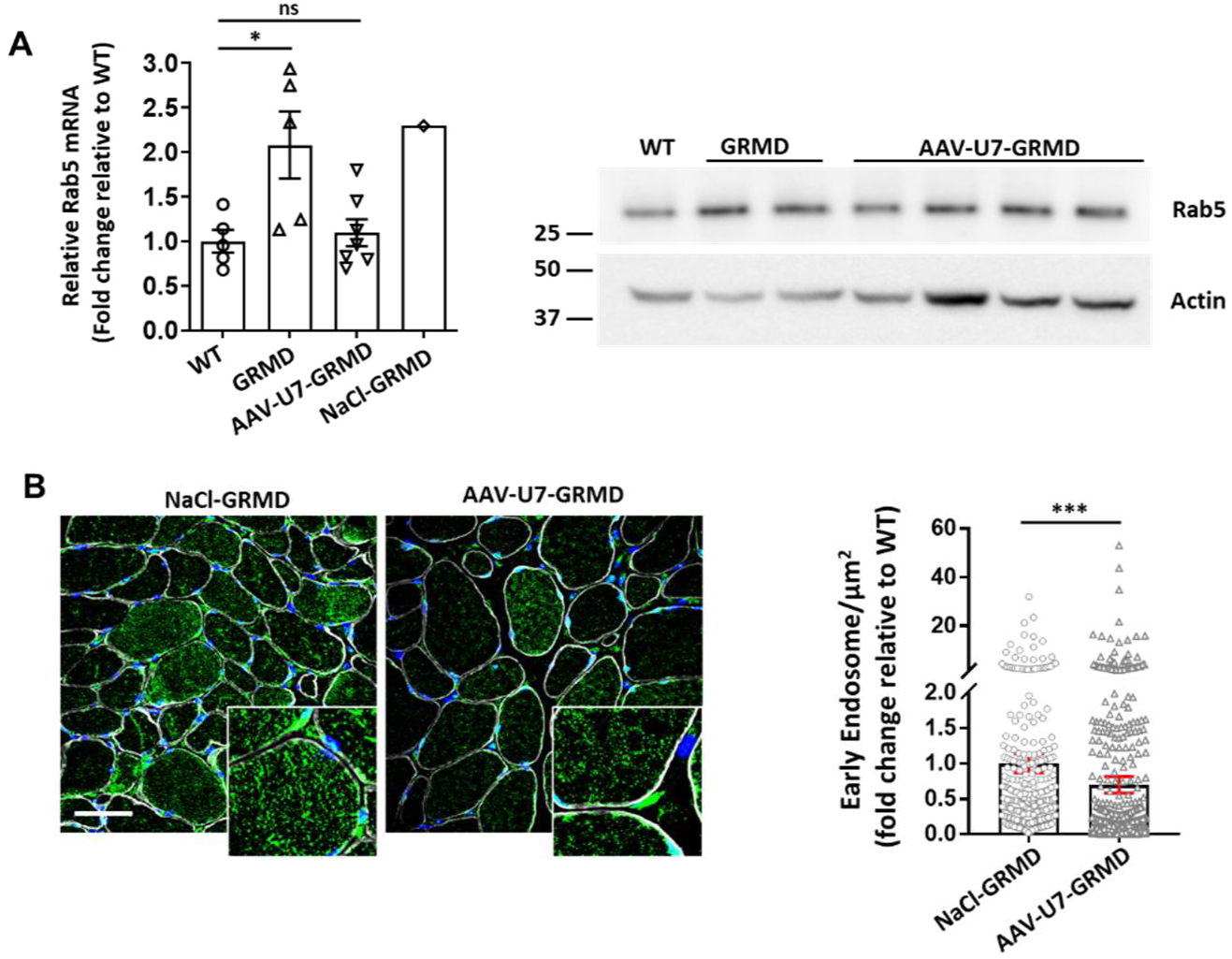
Restored early endosomes number and Rab5 expression in treated GRMD dogs. **(A)** The expression of Rab5 mRNA analyzed by RT-PCR (left) and Rab5 protein analyzed by western blot (right) in muscles of WT dogs (n=5 for mRNA or n=1 for protein), GRMD (n= 5 dogs for mRNA or n=2 dogs for protein), AAV-U7snRNA treated GRMD to restore dystrophin (n = 7 biopsies from 3 treated GRMD for mRNA, NaCl-treated controls (NaCl-GRMD, n=1), or n=2 biopsies per dog from 2 treated GRMD (for protein) showing that expression of Rab5 is decreased by dystrophin restoration in treated GRMD. Actin was used as a loading buffer in western blot. **(B)** Representative sections of muscle biopsies isolated from moderately affected GRMD and AAV-U7snRNA treated GRMD labeled with DAPI to mark nuclei (blue), anti-caveolin antibody to mark the plasma membrane (white) and anti-EEA1 antibody to mark early endosomes (green). Quantification of EEA1 positive puncta showed a significant decrease in early endosome staining in GRMD treated with an AAV-U7snRNA (n=2 biopsies from 1 treated GRMD) compared to NaCl-treated controls (NaCl-GRMD, n=1) showing that the therapeutically restored dystrophin allowed a partial restoration of early endosomes number compared to controls. The data are represented as the mean ± SEM of at least 500 fibers per dog analyzed on confocal images. Sale Bar = 50 µm. Statistics: one-way ANOVA with a post hoc Bonferroni test for panel A (Rab5 mRNA) and unpaired Student’s t-test (two tailed) for panel B (EEs quantification), *p<0.05; ****p<0.0001, ns: non-significant.

## Discussion

The development of new therapeutics and refinement of existing ones for DMD lies in the understanding of the mechanisms disturbing the muscle homeostasis. The absence of expression of the full-length dystrophin in muscles leads to intracellular alterations including severe damage in lysosomal biogenesis and impaired autophagy in human, dog and mouse models of DMD^11,12,42^. Moreover, it triggers major transcriptomic and functional abnormalities in myoblasts^35,36^, cells that play central roles in muscle regeneration and delivery of therapeutic ASO to dystrophic muscles^43^. In accordance with this, we demonstrated an altered acidification of the endosomal-lysosomal pathway associated with a lower degradative capacity in DMD muscle cells.

Dysregulated lysosomal pH was already shown to disrupt DMD muscle homeostasis and muscle regeneration in zebrafish^44^, and it constitutes the primary pathogenic mechanism causing or deteriorating the disease in some cases of neurodegenerative diseases including Alzheimer’s disease (AD) and Parkinson’s disease (PD)^45^. In addition, the lower acidification of endosomal lumen reported here should alter the vesicular hydrolases activities in DMD muscle cells, impacting the sorting and routing of cargo as well as its degradation^16^. The degradative capacity of the endolysosomal pathway would also be impacted if maturation or trafficking of early endosomes to lysosomes is disturbed as previously described in diseases, such as AD and PD^46^. In these pathologies, this defect is characterized by increased number of early endosomes organized in clusters, increased Rab5 GTPase and abnormally increased hydrolases within early endosomes in pyramidal neurons of the cortex^47^. Similarly, we found in DMD myoblasts and myotubes an increase in the number and size of early endosomes, sometimes grouped in clusters, suggesting a disruption of the endosomal trafficking and maturation of early endosomes. This maturation involves concomitant retrograde transport along microtubules (MT) and graded fusion with late endosomes and lysosomes^48^, and a functional endosomal pathway is necessary for lysosomes biogenesis^49–52^. In the current study, the low density of late endosomes and large perinuclear lysosomes concomitant to the high density of early endosomes in DMD muscle cells supports a model in which lysosomes fail to form due to reduced endosomal trafficking. In DMD cells, EE accumulation seems not due to an increased endocytosis suggesting that EE accumulation mainly results from a blockage of the gradual maturation of early endosomes to late endosomes and lysosomes. This could hinder the gradual vesicular pH drop and impact the lysosome and autophagy degradation pathways that are known to be defective in human and animal models of DMD^11,12,42,53,54^.

The first symptoms of DMD appear in early childhood, around 2–3 years of age, with loss of ambulation between 9 and 12 years. In GRMD, limb weakness and exercise intolerance start around 2 to 3 months of age (analogous to ∼3 years of age in humans)^55^. At this age of onset of symptoms, we observed an increase in the abundance of early endosomes at early stages in muscles from DMD patients (3 years of age) and severely affected GRMD dogs (2 months of age). This endosome dysregulation which is maintained at later stages (10 years in patients and 6 months in dogs) showing a long-term alteration affecting different muscle types with different mutations of dystrophin. The 2–6 months of age in GRMD is a period of particularly fast disease progression that corresponds to the 5–10 years period in human DMD patients^55^. We compared the endosomal defect in two subgroups of differentially affected GRMD dogs (moderate and severe) at 2 and 6 months of age. The defect was more pronounced in the severely affected subgroup as early as 2 months, while in the moderately affected subgroup, it became significant at 6 months, correlating with disease severity and progression. This finding raised the possibility that abundance of early endosomes is the consequence of muscle regeneration of the “dystrophic program”. Interestingly, our data show that the localization, density and number of early endosomes are already disturbed in muscle fibers of 2 week-old mdx mice, i.e. before the massive necrosis/regeneration, suggesting that these defects are due to the dystrophin lack *per se*. Given that lysosome formation is defective in mdx skeletal muscle^10,11,42^, our findings reinforce the hypothesis that lysosome impairment could be due, at least in part, to the defect of endosome-dependent lysosome maturation in DMD. Due to the involvement of the decreased lysosomal abundance in autophagic dysfunction in muscles of GRMD and mouse DMD muscles^11,42,56^, we may hypothesize that early endosome dysregulation plays a substantial role in lysosome biogenesis and formation of immature autophagic vesicles resulting in autophagy dysfunctions in DMD.

Defects in early endosomes has been observed in several disease conditions that underly defects in endosome maturation, however, the exact mechanism leading to this phenotype remains unclear. Both actin and microtubule (MT) networks govern the intracellular distribution of the early endosomes and depolymerization of MT with nocodazole resulted in accumulation of enlarged or clustered early endosomes in the periphery of cells^57,58^. Altered production of reactive oxygen species (ROS) via NADPH Oxidase 2 (NOX2) in *mdx* muscles was shown to mediate MT disorganization as well as impairment of both autophagy and lysosomal formation^11,59^. Thus, it is possible that NOX2 ROS dependent MT disorganization impacts early endosomes to lysosomes trafficking and maturation, limiting the degradation of the endolysosomal/autophagy systems. NOX2 ROS is also upregulated early in the progression of the disease (18-19 day old) in mdx^60^, prior to changes in the MT network^8,61–64^. Rab5 GTPase known to be the main regulator of early endosomes homeostasis^65^, leads to abnormal number of endosomes as well as their enlargement when overexpressed^66^. In line with these studies, we found that Rab5 is upregulated in case of dystrophin deficiency as shown in human DMD myoblasts, mdx and GRMD muscles suggesting a role of Rab5 in EE accumulation in DMD. Normalization of Rab5 expression and endosomal abnormalities after dystrophin restoration in GRMD dogs and by Rab5 knock-down in human DMD muscle cells confirms this hypothesis by showing the control of Rab5 by dystrophin expression. This is also supported by the absence of endosome defect in LGMD2C and FSHD1, i.e. two dystrophies not related to dystrophin. Interestingly, Rab5 expression increases has been reported in affected neurons in AD^47,67,68^ as well as in a mouse model of Down syndrome disease^69^ where early endosomal abnormalities hampers degradation of toxic β−amyloid (Aβ) species (Aβ40 and Aβ42). Of note, increased levels of Aβ42 has been involved in cognitive function decline^70^ and muscle degenerative process^71^ in DMD patients suggestive of common pathogenic mechanisms between neurodegenerative diseases and DMD with a potential central role of Rab5 upregulation, EE defects and Aβ42 in the cognitive and muscle impairment in DMD.

In conclusion, our results better define the DMD pathophysiology by highlighting the involvement of Rab5 and early endosomes. Further studies are needed to investigate the potential deleterious effect of this pathophysiological pathway not only in the muscle homeostasis but also in endosomal transport and perinuclear release of therapeutic AAVs and ASOs in DMD. A deeper understanding of these abnormalities will lead to more efficient therapeutic approaches and for a better monitoring of the disease progression.

## Materials and Methods

### Biological resources

*<u>Dog muscle samples</u>* used in the present study (summarized in Supplementary Tables S1) were obtained from the Boisbonne Center for Gene and Therapy (ONIRIS, INSERM, Nantes, France) and the Ecole Nationale Vétérinaire d’Alfort (Maisons-Alfort, France). Biceps femoris muscle biopsies were obtained from 6 untreated GRMD dogs, 3 with a loss of ambulation at the age 6 months (severe form) and 3 still ambulant at this timepoint (moderate form). Muscle biopsies were surgically harvested at 2 and 6 months of age on each of these 6 dogs, as well as on 6 healthy littermates (n=3 at 2 months and n=3 at 6 months), under general anesthesia and proper analgesia (propofol, isoflurane in 100% O2, morphine). Muscle samples from treated GRMD dogs were part of preclinical studies which were based on the delivery of NaCl or therapeutic AAV-U7E6/E8 for specific exon skipping of exons 6 to 8 in order to restore an in-frame dystrophin mRNA as described in the two published preclinical studies^40,41^. Surgical biopsies of the injected dogs were carried out at three^41^ or six^40^ months after injection. Muscle biopsies of healthy dogs, untreated- and treated-GRMD were snap-frozen in isopentane cooled in liquid nitrogen for immunohistology or flash frozen in liquid nitrogen for western-blot and RT-PCR.

All procedures were approved by the Ethical Committee of EnvA, ANSES, and UPEC and by the French Ministry of Research in accordance with the relevant guidelines and regulations under the approval number 20/12/12-18. The Institutional Animal Care and Use Committee of the Region des Pays de la Loire (University of Angers, France) approved the protocol.

*<u>Biopsies</u>* from three healthy individuals and three DMD patients were included in the present study (Supplementary Table S2). Skeletal muscle sections of 10µm were obtained from the Morphological Unit of the Institute of Myology (Paris, France) taken for diagnostic purposes^72^

*<u>Immortalized skeletal muscle cell lines</u>* derived from healthy donors (controls, Ctrl) and from DMD (deletions of exons 45–52 or exons 52), LGMD2C (Limb-Girdle Muscular Dystrophy, Type 2C) and FSHD (facioscapulohumeral muscular dystrophy) patients as well as non-immortalized muscle cells were obtained from the MyoLine platform of the Myology Institute (Paris, France). Immortalized muscle cells were generated as previously described^73^.

*<u>For mouse muscles</u>,* wild-type (WT) and mdx mice were anesthetized by intraperitoneal injection of ketamin/xylazine and sacrified by cervical dislocation. The extensor digitorum longus (EDL) from 2-week-old or 3-months-old mice or Tibialis Anterior (TA) muscles from 3-months-old mice were surgically isolated for fiber isolation or for western-blot.

### Cell culture

Immortalized skeletal muscle cell lines and non-immortalized ones were maintained at 37 °C and 5% CO_2_ in a proliferation medium composed of 4 volumes of DMEM-High Glucose medium for 1 volume of medium 199 (Life Technologies, Villebon-sur-Yvette, France) supplemented with 5 μg/ml of insulin (Sigma-Aldrich Chimie, Saint Quentin Fallavier, France), 5 ng/ml of hEGF (Life Technologies), 0.5 ng/ml of bFGF (Life Technologies, Villebon-sur-Yvette, France), 0.2 μg/ml of dexamethasone (Sigma-Aldrich Chimie, Saint Quentin Fallavier, France), 25 μg/ml of fetuin (Life Technologies, Villebon-sur-Yvette, France), 20% of fetal bovine serum (Life Technologies). Differentiation was induced at confluence by replacing the growth medium with DMEM supplemented with 0.1 ng/ml gentamycine and 10 μg/ml of insulin (Sigma-Aldrich Chimie, Saint Quentin Fallavier, France).

### siRNA transfection

Muscle cells were seeded at 60,000 per well in 24 well plates 24h before transfection. Specific knockdown of Rab5 was obtained by transfection of siRNA (siRab5) (Silencer Select siRNA, Life Technologies, Villebon-sur-Yvette, France) at a final concentration of 6 pM using Lipofectamine RNAiMAX (Life Technologies SAS, Villebon-sur-Yvette, France) according to the manufacturer’s protocol for adherent cells. In these experiments, Silencer Select Negative Control n°2 (Life Technologies) was used as a control (siCtrl).

### Protein lysates and western-blot

Human muscle cells and animal tissues were lysed in RIPA buffer (150 mM NaCl, 50 mM 4-(2-hydroxyethyl)-1-piperazineethanesulfonic acid, pH 7.4, 5 mM ethylenediamine tetra acetic acid, 1% NP-40, 0.5% sodium deoxy-cholate, 0.1% sodium dodecyl sulfate, 1 mM PMSF containing 1% of a protease inhibitor cocktail (Sigma-Aldrich Chimie, Saint Quentin Fallavier, France), and centrifuged 20 min 11000g at 4°C. Protein extracts (60 μg) were denaturated in Laemmli buffer 2X added of 10% of 2-mercaptoethanol 2 min at 96°C. Proteins were resolved by SDS–PAGE (4–12%, Life Technologies, Villebon-sur-Yvette, France) and transferred to PVDF. Membranes were blocked in Tris-buffered saline (TBS) 0.1% Tween-20 with 5% non-fat dry milk 1h at RT and incubated overnight at 4°C with mouse monoclonal antibody against Rab5 (1/500; Santa Cruz Biotechnology, Heidelberg, Germany), with mouse antibody against LAMP1 (1/1000; Santa Cruz Biotechnology, Heidelberg, Germany) or with rabbit monoclonal Actin antibody (1/1000; Sigma-Aldrich Chimie, Saint Quentin Fallavier, France). After being washed in TBS 0.1% Tween-20, membranes were incubated for 1h at RT with secondary antibodies: goat anti-rabbit-horseradish peroxidase (HRP) (1/10000), sheep anti-mouse HRP (1/10000) or Goat anti-rabbit HRP (1/10000) (Jackson Immunoresearch Inc. West Grove USA). Western-blots were revealed by Pierce™ ECL Western-Blot Substrate (Life Technologies) with ChemiDoc MP Imaging System (Bio-Rad Laboratories, Marnes-La-Coquette, France).

### Transcript quantification

For mRNA quantifications, total RNA was isolated from cultured muscle cells using NucleoSpin RNA Virus (Macherey-Nagel SAS, Hoerdt, France) and from pooled muscle sections of animal biopsies using TRIZOL-reagent (Life Technologies, Villebon-sur-Yvette, France). Reverse transcription was performed on 200 ng of RNA by using the High-Capacity cDNA Reverse Transcription Kits (Life Technologies). Rab5 transcripts were quantified and normalized to mRNA levels of beta-2-microglobulin (β2M) for human muscle cell cultures, ribosomal protein L32 (RPL32) for dogs and 18S ribosomal RNA (Rn18s) for mice on Quant Studio 3 (Applied Biosystems, Life Technologies) using Master Mix TaqMan Fast Advanced (Life Technologies). The TaqMan Gene Expression Assays (Life Technologies) were: human Rab5 (ID : Hs00702360_s1), mouse Rab5 (ID : Mm00727887_s1), dog Rab5 (ID : Cf00991291_m1), B2M (ID: Hs00984230_m1), Rpl32 (ID : Cf03986503_g1) and Rn18s (ID : Mm03928990_g1), used under the conditions: 50°C for 2 min, 95°C for 10 min, 40 cycles of 15 s at 96°C and 1min at 60°C.

### EFG uptake

Cells were cultured in DMEM, 0.02M HEPES, 1% BSA without FCS during 45 min at 37°C in 24-well plate and were exposed to DMEM, 0.02M HEPES, 1% BSA supplemented with 25 µg/ml of Alexa 488-tagged EGF (Alexa-488-EGF) or pHrodo-tagged EGF (pHrodo-EGF) ice for 1h, washed with iced Live Cell Imaging Solution (Life Technologies, Villebon-sur-Yvette, France) and incubated at 37°C in the microscope thermostatic chamber for up to 60min in DMEM, 0.02M HEPES, 1% BSA for live imaging. Fluorescent-positive surface was quantified on stack projection using ImageJ software and normalized to the total cell surface.

### Transferrin uptake (supplementary Fig.S4)

Cells were cultured in DMEM, 0.02 M Hepes, 1% BSA without FCS during 45 min at 37°C in 24-well plate containing glass coverslips and were exposed to DMEM, 0.02 M Hepes, 1% BSA supplemented with 25 µg/ml of Alexa-Fluor488-Transferrin (Life Technologies, Villebon-sur-Yvette, France) or biotinylated-transferrin on ice for 1h and, washed with iced Live Cell Imaging Solution (Life Technologies) for Alexa-Fluor488-Transferrin (Life Technologies) or PBS for Biot-Tfn and incubated at 37°C in DMEM, 0.02 M Hepes, 1% BSA. Cells were washed 3 times with PBS at each time point and fixed in 4% paraformaldehyde at room temperature for 15 min.

For the biotinylated-transferrin, cells were lysed in RIPA buffer and proteins were submitted to western-blot. Biotin was detected using horseradish peroxidase (HRP)-conjugated streptavidin (Pierce, Life Technologies, Villebon-sur-Yvette, France) and the Supersignal West PicoChemiluminescent kit (Life Technologies). For Alexa-Fluor488-Transferrin, stacks of cell images (0.2 μm interval) were obtained using a Leica SP2 confocal microscope. Fluorescent-positive surface was quantified on stack projection using ImageJ software and normalized to the total cell surface.

### Immunostaining

*<u>For human myoblasts and myotubes</u>*, cells were grown on coverslips, washed with PBS, and fixed with 4% paraformaldehyde in phosphate-buffered saline (PBS) for 10 min. Cells were permeabilized with 0.5% Triton X-100 (Sigma-Aldrich Chimie, Saint Quentin Fallavier, France), and non-specific signals were blocked in PBS supplemented with 4% bovine serum albumin (BSA) for 1h at room temperature (RT). Cells were then incubated with a goat monoclonal antibody EEA1 (1/500; SICGEN, Cantanhede, Portugal), a rabbit monoclonal antibody Rab7 (1/200; Santa Cruz Biotechnology, Heidelberg, Germany) and a mouse monoclonal antibody LAMP1 (1/100; Santa Cruz Biotechnology, Heidelberg, Germany) in PBS supplemented with 0.5% Triton X-100 for 2h at room temperature, washed with PBS and incubated with the appropriate secondary antibodies labeled with Alexa Fluor 488 or Alexa Fluor 568 (1/1000; Life Technologies, Villebon-sur-Yvette, France) for 45 min at RT and, with 4’,6-diamidino-2-phenylindole (DAPI) (Sigma-Aldrich Chimie, Saint Quentin Fallavier, France) for 5 min for nuclei staining. Cells were then washed in PBS and slides were mounted in Fluoromount mounting medium (Southern Biotech, Birmingham, USA) for the microscopy analysis.

*<u>For mouse myofibers</u>,* the extensor digitorum longus (EDL) or Tibialis Anterior (TA) muscles were surgically isolated for fiber isolation or for western-blot. EDL muscles were dissected from 2-week-old or 3-months-old wild-type and mdx mice and digested in DMEM 0.2% collagenase 1h at 37°C. Fibers were separated mechanically and fixed in 4% paraformaldehyde for 15min. Myofibers were fixed and permeabilized as described above, blocked in PBS supplemented with 0.7% Triton X-100, 0.1% Tween-20 and 4% BSA overnight (ON) and then incubated ON with a mouse monoclonal antibody against EEA1 (1/150; Santa Cruz Biotechnology, Heidelberg, Germany), and a mouse monoclonal antibody against Alpha Tubulin (1/1000; Sigma-Aldrich Chimie, Saint Quentin Fallavier, France). Myofibers were washed, incubated with appropriates secondary antibodies and DAPI and mounted as described above.

*<u>For dog and human muscle biopsies</u>*, cryosections of 8 µm (for dogs) or 10 µm (for human) were fixed in 4% paraformaldehyde for 10 min, washed in PBS, permeabilized with 0.25% Triton X-100 at RT and blocked in PBS supplemented with 10% fetal bovine serum (FBS) for 1h. Sections were incubated with a goat monoclonal antibody against EEA1 (1/100; SICGEN, Cantanhede, Portugal) and Caveolin 3 (1/750, BD Biosciences, Pont de Claix, France) antibodies (for dog sections) or EEA1 and a rabbit monoclonal antibody laminin alpha-2 (Abcam limited, Cambridge, UK) antibodies (for human sections) in PBS supplemented with 2% FBS ON at 4°C and washed with PBS. Sections were then incubated for 1h with appropriate secondary antibodies in same buffer, washed with PBS, incubated with DAPI and mounted in Fluoromount-G (Cliniscience, Amsterdam, Netherlands).

### Microscopy and imaging

Confocal images were acquired using a Nikon Ti2 microscope, driven by Metamorph (Molecular Devices), equipped with a motorized stage and a Yokogawa CSU-W1 spinning disk head coupled with a Prime 95 sCMOS camera (Photometrics) equipped with a 100x oil-immersion objective lense.. DAPI, Alexa Fluor 488, Alexa Fluor 568 and Alexa Fluor 647 were sequentially excited. Z-series from the top to the bottom of cells were sequentially collected for each channel with a step of 0.1-0.3 μm between each frame. Image quantification was performed using National Institutes of Health’s FIJI. The number and size of endosomes was determined using the Cell Profiler Software or the 3D Objects Counter function of Fiji software.

Widefield fluorescence microscopy was performed using a Nikon Ti2 microscope equipped with a motorized stage and coupled with CMOS DS-Ri2 Nikon camera at 10x/0.45 NA objective and a Lumencor LED fluorescent lamp was used. The images were acquired using NIS 5.11 software with RGB 8-bit in 5 × 5 slide Scan mode at a pixel resolution of 0.73 μm/px. For muscle sections, fibers were segmented with CellPose using NVIDIA GeForce RTX 3080 Ti GPU and quantified with QuPath.

### Electron microscopy

Cells were grown on coverslips, washed with PBS, fixed in phosphate buffer 0.1 M, 4% paraformaldehyde and 0.05% glutaraldehyde. After aldehyde masking with PBS supplemented with 0.1 M glycine, cells were blocked in PBS supplemented with 5% goat serum, 5% BSA, then incubated with a rabbit antibody against EEA1 (Cell Signaling Technology, Inc. Danvers, USA) and finally with nanogold-labeled goat against rabbit antibody (Aurion, Netherlands). This step was followed with a silver amplification (HQ silver, Nanoprobes, NY). After 15 min post-fixation in 1% OsO4, sections were dehydrated in graded acetone and finally embedded in Epon resin. Ultrathin sections were lightly post-stained in uranyl acetate and lead citrate, examined using a Philips CM120 electron microscope operated at 80kV and imaged with a SIS Morada digital camera.

## Acknowledgements

We thank Zoheir Guesmia from MyoImage platform for image analysis, Christel Gentil and Stéphanie Duguez for technical advices, Anne Bigot and Vincent Mouly from MyoLine platform for providing primary and immortalized human muscle cells, the staff of the animal facilities of Sorbonne University (CEF, UMS28, Pitié-Salpêtrière) for the handling and care of mice, the Centre d’Elevage du Domaine des Souches, the team of the neurobiology laboratory in the Alfort veterinary school as well as Boisbonne Center for Gene Therapy (ONIRIS, INSERM, Nantes, France) for the handling and care of GRMD dog colonies. We thank Adeline Vulin for providing muscle biopsies of AAV1-U7- and NaCl-treated GRMD from the published preclinical study ^40^. This work was supported by Duchenne Parent Project NL (DPP-NL), the Association Institut de Myologie (AIM), Association AFM-Téléthon, the Institut National de la Santé et de la Recherche Médicale (INSERM) and Sorbonne University.

