## Supplemental data for "Early endosome disturbance and endolysosomal pathway dysfunction in Duchenne muscular dystrophy"

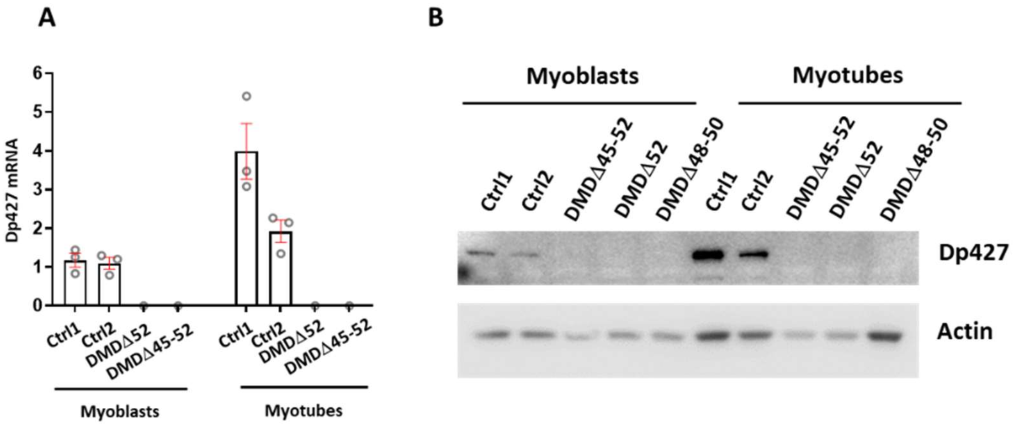


**Supplementary Fig. S1: Dystrophin (Dp427) expression in undifferentiated and differentiated muscle cells derived from human biopsies.**

**(A)** Real time RT-PCR and **(B)** Western blotting analysis of dystrophin (Dp427) in proliferating/undifferentiated (myoblasts) and differentiated (myotubes) muscle cells derived from 2 healthy controls (Ctrl1 and Ctrl2) and 3 DMD patients with 3 distinct deletions (DMDΔ45-52, DMDΔ52 and DMDΔ48-50) showing low levels of Dp427 mRNA **(A)** and protein **(B)** expression in control myoblasts compared to control myotubes (Ctrl1 and Ctrl2). No expression was observed in DMD myoblasts and myotubes. Transcripts were quantified using the Power SYBER Green PCR Master mix (Thermofisher Scientific) and the following primers: dysE4-E5 primers (F: GGCACTGCGGGTCTTACA and R: CATCCACTATGTCAGTGCTTCCTAT) were used to amplify dystrophin transcripts and the ribosomal phosphoprotein (RPLO) mRNA amplification was used for normalization (F: CTCCAAGCAGATGCAGCAGA and R: ATAGCCTTGCGCATCATGGT). Mandra1 antibody (Developmental Studies Hybridoma Bank) that recognizes the end of the C-terminal domain of dystrophin and actin antibody (sigma) was used in Western-blot.

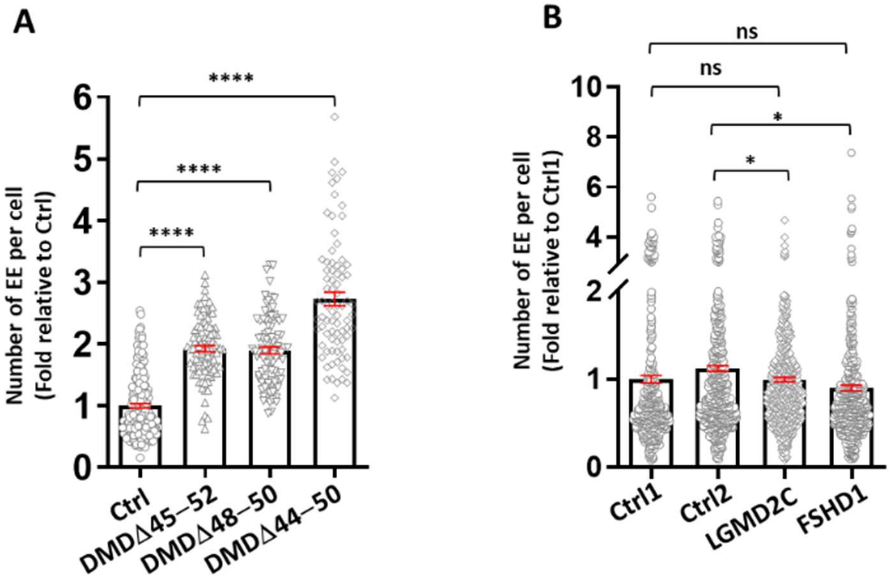


**Supplementary Fig. S2: Analysis of early endosomes in muscle cells of DMD, LGMD2C and FSHD1 patients.**

**(A)** Quantification of early endosomes (EEs) after EEA1 immunostaining showed an increased number of EEA1 positive puncta in non-immortalized DMD primary myoblasts (DMDΔ45-52 and DMDΔ48-50, DMDΔ44-50) compared to control myoblasts. **(B)** Quantification of early endosome positive puncta after EEA1 immunostaining in LGMD2C and FSHD1 myoblasts derived from human biopsies compared to 2 control cell lines (Ctrl1 and Ctrl2). The data are represented as the mean ± SEM of 4 independent experiments analyzed on confocal images (at least 80 cells analyzed). Statistics: one-way ANOVA with a post hoc Bonferroni test, *p<0.05, ****p<0.0001, ns : non-significant

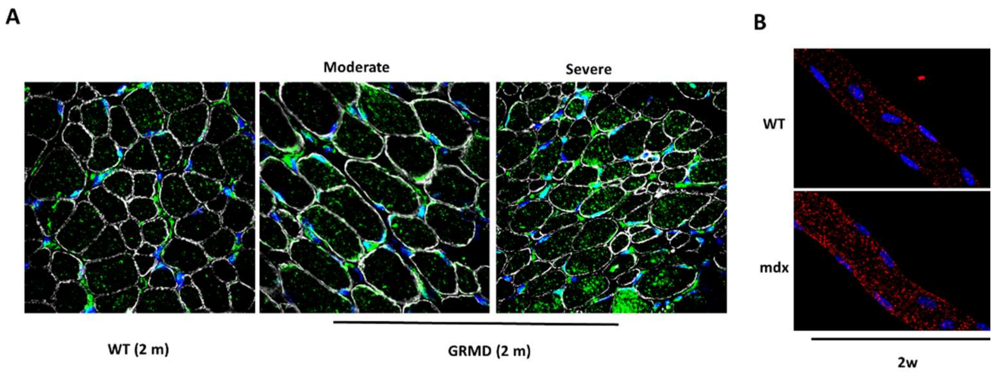


**Supplementary Fig. S3: Distribution of early endosomes in muscle fibers from GRMD and mdx**

Representative confocal images of **(A)** sections of muscle biopsies isolated from severely or moderately affected GRMD and healthy control at 2 months of age labeled with Dapi to mark nuclei (blue), anti-caveolin antibody to mark the plasma membrane (white), and anti-EEA1 antibody to mark early endosomes (green) **(B)** myofibers isolated from EDL muscles of wild type (WT) and mdx at 2 weeks of age labeled with Dapi to mark nuclei (blue) and anti-EEA1 antibody to mark early endosomes (red).

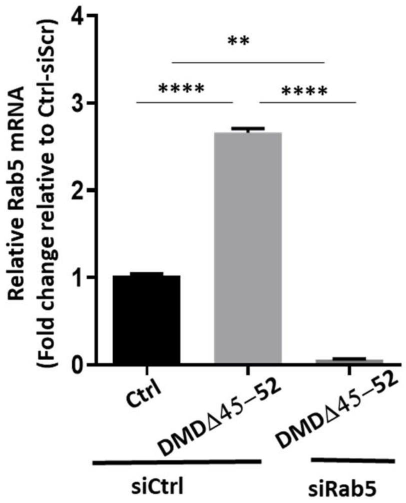


**Supplementary Fig. S4: Rab5 siRNA efficiency**

Real time RT-PCR analysis of Rab5 mRNA expression showing the efficacy of Rab5 silencing in DMD myoblasts transfected with siRNA directed against Rab5 (siRab5) compared to DMD and control myoblasts (Ctrl) treated with a control siRNA (siCtrl). The data are represented as the mean ± SEM of

3 independent experiments. Statistics: one-way ANOVA with a post hoc Bonferroni test, **p< 0.01, ****p<0.0001.

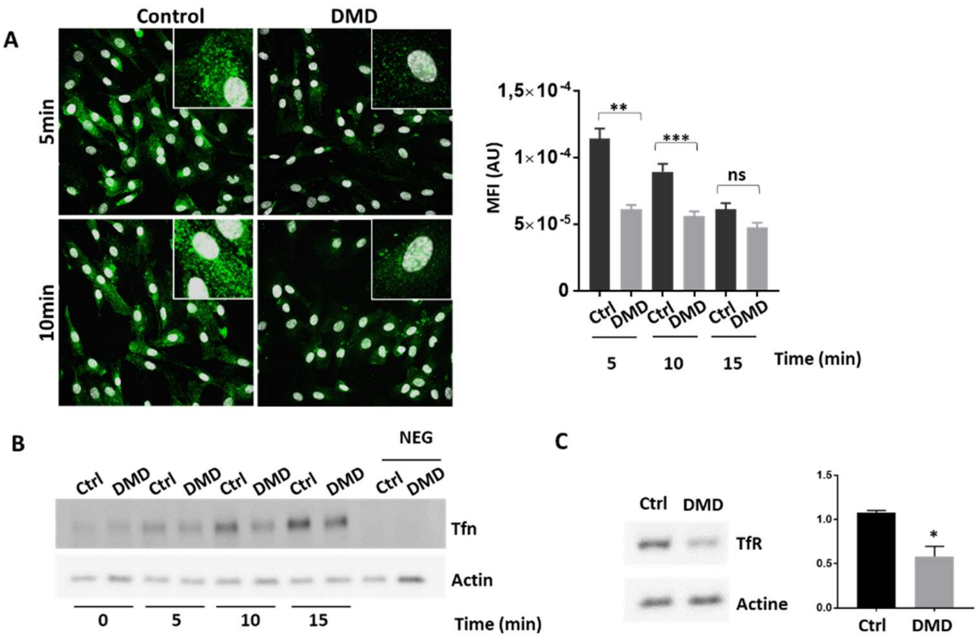


**Supplementary Fig. S5: Endocytosis in DMD Δ45-52 and control myoblasts.**

**(A)** Representative images (left) of 5- and 10-min uptake of Alexa Fluor 488-Tfn (green) and quantification of the mean fluorescence intensity (MFI) per cell area (right) of 5- 10- and 15 min uptake in DMD and control myoblasts. The data are represented as the mean ± SEM of 4 independent experiments on confocal images (at least 200 cells per experiment). **(B)** Western blot of Biot-Tfn illustrating the kinetic of Biot-Tfn uptake from 0 to 15 min in DMD and control myoblasts (n=4). **(C)** Western blot of the transferrin receptor (TfR) showing a two-fold decreased expression of the TfR in DMD cells compared to control (n=3). Actin was used for normalization of western blot (n=3). Statistics: one-way ANOVA with a post hoc Bonferroni test for Alexa-488-Tfn and unpaired Student's t-test (two tailed) for TfR western-blot, *p<0.05, **p<0.01, ***p<0.001, ns : non-significant

**Supplementary Table S1: Dog muscle samples used in this study**

| Group | Dog | Injection | Sample collection | Muscle type | WB | RT-qPCR | Immunostainings anti-EEA1 | Microscopy analysis |
|---|---|---|---|---|---|---|---|---|
| Healthy controls | C1 | / | 2 months | Biceps femoris | / | / | X Fig. S3 | X |
| | C2 | | | | / | / | X | X |
| | C3 | | | | / | / | X | X |
| GRMD moderate | D1 | | | | / | / | X Fig. S3 | X |
| | D2 | | | | / | / | X | X |
| | D3 | | | | / | / | X | X |
| GRMD severe | D4 | | | | / | / | X Fig. S3 | X |
| | D5 | | | | / | / | X | X |
| | D6 | | | | / | / | X | X |
| Healthy controls | C4 | / | 6 months | | / | X Fig. 6 and Fig. 7 | X Fig. 5 | X |
| | C5 | | | | / | | X | X |
| | C6 | | | | / | | X | X |
| GRMD moderate | D1 | | | | / | / | X Fig. 5 | X |
| | D2 | | | | / | / | X | X |
| | D3 | | | | / | / | X | X |
| GRMD severe | D4 | | | | / | X Fig. 6 and Fig. 7 | X Fig. 5 | X |
| | D5 | | | | / | | X | X |
| | D6 | | | | / | | X | X |
| Healthy | C7 | / | 6 months | Flexor Digitorum Superficialis | / | X Fig. 6 and Fig. 7 | / | / |
| | C8 | | | Biceps femoris | X Fig. 6 and Fig. 7 | | / | / |
| GRMD | D7 | | | Flexor carpi ulnaris | | | / | / |
| | D8 | | | | | | / | / |
| | D8 | | | Extensor carpi radialis | / | | / | / |
| AAV8-U7-E6/E8 treated GRMD (Leguiner C. *et al* , 2014) | D9 | 3 months | 6 months | Flexor Digitorum Superficialis | X Fig. 7 | X Fig. 7 | / | / |
| | D9 | | | Deltoid | | | / | / |
| | D10 | | | Flexor digitorum profundus | | | / | / |
| | D10 | | | Supinator | | | / | / |
| AAV1-U7-E6/E8 treated GRMD (Vulin A. *et al* , 2012) | D11 | 14 months | 20 months | Biceps femoris | / | | / | X |
| | D11 | | | | / | | / | X |
| | D11 | | | | / | | X Fig. 7 | X |
| NaCl treated GRMD | D12 | | | | / | | | X |

GRMD, golden retriever muscular dystrophy

X, used in the RT-qPCR or WB or microscopy analysis

**Supplementary Table S2:** Human samples used in this study

| Group | Deletion | Age (year-old) | Muscle type |
|---|---|---|---|
| Healthy | / | 3 | Deltoïd |
| | | 5 | |
| | | 10 | |
| DMD | Delta 45-52 | 4 | |
| | Delta 48-50 | 10 | |
| | Delta 52 | 5 | Quadriceps |

DMD, Duchenne Muscular Dystrophy
